# From Fear to Feast: Rattlesnakes Navigate the Landscape of Fear to Optimize Foraging

**DOI:** 10.1101/2024.11.14.623629

**Authors:** Oceane Da Cunha, Rio P Dominguez, L. Miles Horne, Joshua J. Mead, Corentin Fournier, Jerry D Johnson, Brett M Seymoure

## Abstract

According to optimal foraging theory, mesopredators should forage in areas where their prey is abundant while avoiding high predation risk. Here, we investigate how environmental factors influence mesopredators’ abilities to minimize spatiotemporal overlap with predators while increasing spatiotemporal overlap with prey. We paired thirty western diamond-backed rattlesnake (*Crotalus atrox*) 3D-printed replicas with game cameras in West Texas for two years to quantify several spatiotemporal factors affecting prey availability and predation risk. Concurrently, 25 *Crotalus atrox* were radiotracked at the same site to gather activity and microhabitat selection data regarding free-ranging individuals. Random forest algorithms were trained using data obtained from the game camera and applied to predict the probability of predation and the probability of prey encounter for each radiotracking event. Time of day, month, vegetation structure, and concealment percentage, all had a significant association with the probability of predation and the probability of prey encounter. Our results suggest that rattlesnakes choose to be active when and where the probability of prey encounter was significantly higher than the probability of detection by predators, thus following optimal foraging theory. Our results demonstrate that mesopredators increase chances of prey capture while reducing predator detection in natural setting.

## Introduction

Predator-prey interactions play a key role in the survivorship of an individual. As such, predator-prey interaction studies have been foundational for understanding broad ecological themes such as community structure (Kneitel and Chase 2004), trophic cascades (Mooney et al. 2010), and biodiversity (Letnic et al. 2012). Overall, predator-prey interactions are pivotal in shaping ecosystem structure and function (Berger et al. 2001; Hawlena and Schmitz 2010). Environmental factors such as temperature and habitat complexity have been noted to influence predator-prey interactions (Wasserman et al. 2016). Moreover, the dynamics between predators and prey are influenced not only by behaviors during interactions (McGhee et al. 2013; Belgrad and Griffen 2016), but also by the behaviors of both prey and predators apart from direct interactions.

Ideally, prey should minimize interactions with their predators. Indeed, many prey modify their morphology, physiology, and behavior in response to predation risk (Lima 2009; Cresswell 2011; Kishida et al. 2014), which is the probability of an individual to be predated by another organism (Pettorelli et al. 2015). For instance, changes in habitat use (Creel et al. 2005), foraging strategy (Winnie and Creel 2007), and movement patterns (Fortin et al. 2005) have been observed in response to predation risk. Such behavioral modifications are costly for prey species and have been found to negatively impact prey growth (Pangle et al. 2007), development (Skelly and Werner 1990), and fecundity (Peckarsky et al. 1993). The indirect effects of predation, such as predation risk, have a significant influence on prey dynamics, sometimes even more prominent than the direct effects of predation (Nelson et al. 2004; Cresswell 2011). Predation risk not only directly affects prey survival but also has the potential to alter community structure. For example, prey will shift their distribution and foraging activity in response to perceived predation risk (i.e. “fear”) thus modifying interspecific prey competition (Lima 1998) with cascading effects across trophic levels (Grabowski and Kimbro 2005). The landscape of fear hypothesis proposes a model to explain these effects and how they cascade from individuals to ecosystems (Brown and Kotler 2004). This model measures how animals perceive their environment on a spatiotemporal scale based on trade-offs between perceived predation risk and activity patterns, thus creating a map of fear across the physical landscape (Bleicher 2017). These landscapes of fear continuously change and are shaped by a variety of biological, ecological, and evolutionary factors (Bleicher 2017; Gaynor et al. 2019).

The landscape of fear does not equally affect the foraging decisions of all species. For example, mesopredators are an interesting group to study in the context of landscape of fear as their foraging decisions are influenced by apex predators (Haswell et al. 2018) and by prey availability (Brown and Kotler 2004). Mesopredators have a central role in predator-prey interactions by serving as both prey and predators within food webs (Mukherjee et al. 2009). Optimal foraging theory suggests that animals must decide what to eat, where to eat, how much time to spend in resources patches, and how to move between resources patches when foraging (Pyke et al. 1977). Following from the optimal foraging theory, specifically optimal patch choice, mesopredators should forage when and where their prey are abundant while avoiding high predation risk (Emlen 1966; MacArthur and Pianka 1966; Pyke et al. 1977). However, studies investigating the foraging decisions of mesopredators in a context of predator-prey interactions are lacking, especially in natural settings. This disparity is partially due to the technical difficulty to observe instances of predation in the wild, particularly for cryptic mesopredators such as rattlesnakes. Nevertheless, rattlesnakes offer a good model to study predator-prey interactions as they are widespread and abundant mesopredators in many North American ecosystems. Due to their ambushing foraging strategy, rattlesnakes spend extended periods of time exposed to predators (Klauber 1956). Thus, rattlesnakes and other mesopredators should face heavy selection pressures to forage when and where prey are in high abundance and predators are in low abundance. However, these pressures are not well understood due to the lack of information regarding predation of rattlesnakes, such as predator encounter rates (Maag and Clark 2022), and other mesopredators. To quantify optimal patch choice in mesopredators, information about predation risk and prey encounter needs to be gathered. Scientists have used rattlesnake replicas made of soft and malleable materials to gather information about predation (e.g. Farallo and Forstner 2012; Harmel et al. 2020). While these types of replicas provide valuable insights, they present some disadvantages such as being time consuming to produce (Behm et al. 2018), being primarily unrealistic, and having limitations about the information gathered from predation events. With the recent development of 3D-printing, replicas can be more biologically accurate and easy to build and manipulate (Bulté et al. 2018). The technology of 3D-printing offers opportunities to ask a wide array of behavioral questions (Igic et al. 2015), including reproduction, foraging, and predation behavior (Behm et al. 2018). While 3D-printed replicas have been slowly introduced in ecology, they have not been extensively used to study predation. Through combining the use of biologically 3D-printed replicas with other technologies, such as radiotelemetry and camera trapping, the trade-offs that mesopredators face involving foraging decisions can finally be quantified.

Here, we aimed to investigate the foraging decisions of the western diamond-backed rattlesnake (*Crotalus atrox*) in natural settings by combining the use of 3D-printed biologically accurate snake replicas, camera trapping, and telemetry data. The main goal of this study was to determine how the landscape of fear affects the foraging decisions of *Crotalus atrox* by looking at factors influencing detection risk and prey availability. Predation risk and prey availability were hypothesized to influence daily activity patterns and micro-habitat selection with the prediction that *Crotalus atrox* are active when and where prey availability is high, and predation risk is low following the optimal foraging theory. To answer these questions, 3D-printed snake replicas associated with game cameras were deployed in the field to assess detection risk and prey availability. Concurrently, western diamond-backed rattlesnakes were radio-tracked and the telemetry data obtained were compared to game camera observations using machine learning algorithms to investigate their foraging decisions. Testing whether these basic ecological predictive approaches work has implications for understanding the long-term evolutionary pressures mesopredators face and how they respond under everchanging environments.

## Material and methods

### Rattlesnake 3D-printed replica design

A preserved specimen of *Crotalus atrox* from the UTEP Biodiversity Collection (Catalogue #: 12333) was scanned with a modified Xbox Kinect^©^ camera (Microsoft, Redmond, VA). This specimen was a male, had a snout-vent length of 77.1 cm and a tail length of 6.3 cm. The replica’s shape (see figure 1) was selected to be in between a coiled and elongated shape to encompass all behavior observed while radiotracking (see radiotracking method). The digital model created from the scan was then adjusted and printed with polylactic acid (PLA) using UltiMaker 3D-printers (UltiMaker, Utrecht, Netherlands). Replicas were printed in two different sizes: a smaller size (total length of 34.1 cm) to represent a juvenile *C. atrox* and a larger size (total length of 85.5 cm) to represent an adult *C. atrox*. The average total length for the radiotracked *Crotalus atrox* in this study was 87.7 cm, thus approximately matching the length of the replicas. Replicas were then painted according to the coloration and pattern of a western diamond-backed rattlesnake. To do so, the body coloration of a wild-caught *C. atrox* was measured in the visible spectrum by placing it in a bucket with white, black, and grey (18% reflectance) standards to calibrate measurements. Pictures were taken with a Canon EOS REBEL T3i camera with a Canon EF 50mm f/1.4 USM lens. To control for light exposure, the bucket was illuminated by a light bulb presenting a spectrum close to natural sunlight (Exo Terra, Baie d’Urfé, CA, Quebec, Exo Terra^©^ halogen basking spot). Images were processed using Image J (Schneider et al. 2012) with the package micaToolbox (Troscianko and Stevens 2015). Coloration was quantified for the dorsum, diamonds, white outline of the diamonds, rattle, and black and white tail bands. Different paint mixes were applied to 3D-printed replicas and colors were measured with the same set-up as previously described. Paint mixes were considered satisfactory for each body part when presenting a significant amount of overlap for blue, red, and green (see supplementary material, figure S1). Then, paint (Acrylicos Vallejo, Barcelona, Spain) was applied to the replica using a combination of airbrushing and paint brushing (see supplementary material, Table S1). Snake scale pattern was created using fishnet stockings and a layer of varnish was applied to protect the paint from rain and sun damage (Liquitex professional matte varnish, Cincinnati, Ohio, USA). Pieces of wood (1 cm x 0.5 cm x 0.5 cm) were glued under the replica to limit their contact to the ground and protect them as ground temperature is known to reach over 60°C during the hottest part of the year.

**Figure 1:**
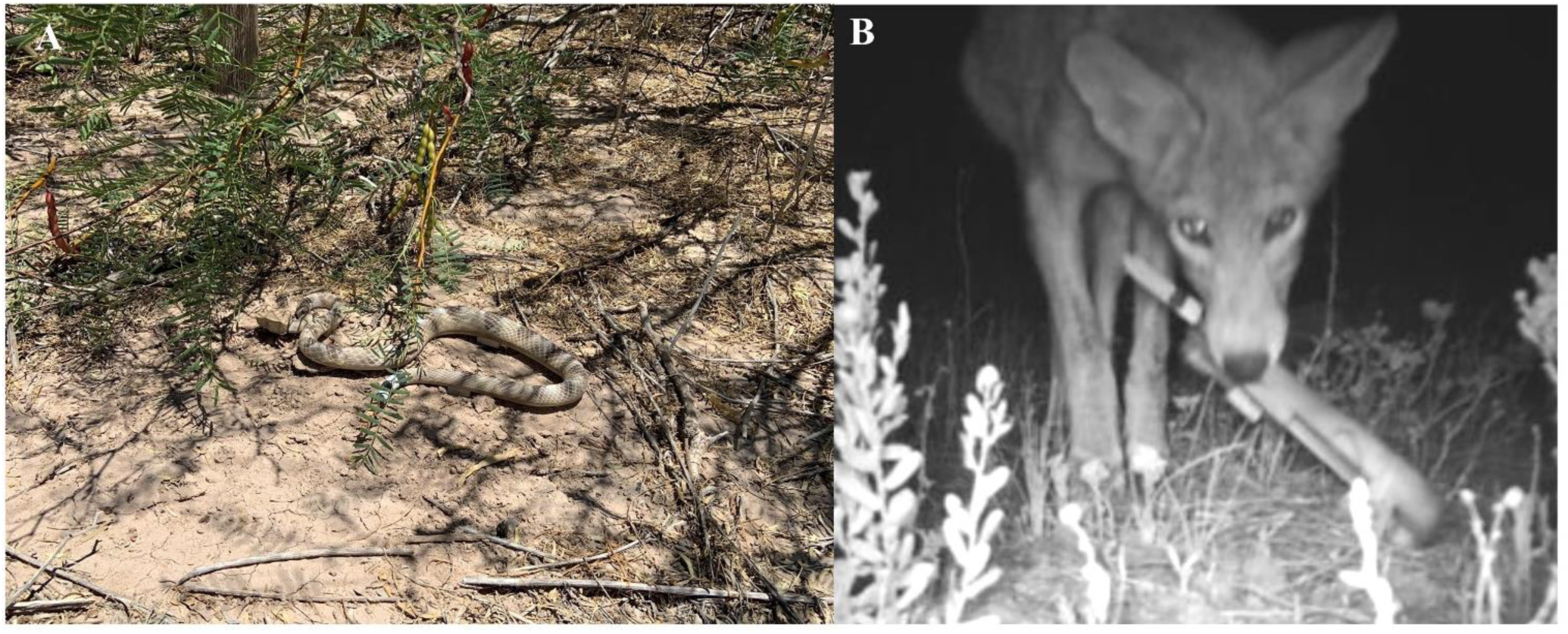
**A**: A rattlesnake 3D-printed replica deployed in the field. **B**: A coyote (*Canis latrans*) predation observation.

### Replica deployment

The replicas were deployed at the Indio Mountains Research Station (IMRS) from June 2020 to December 2022 (figure 1). IMRS is a 161 km^2^ research facility located in Hudspeth County (TX, USA) owned by the University of Texas at El Paso (UTEP). IMRS is characterized by a mixed desert scrub plant community located within the Chihuahuan Desert (Worthington et al. 2022). Remnant earthen cattle tanks can be found within the property from past ranching activities. Tanks constitute an important landscape feature as they still collect water during the rainy season resulting in biotic resource hot spots (DeSantis et al. 2019). For this reason, two main habitats were identified for the purpose of this study: earthen tanks and desert scrub. The replicas were distributed randomly through the landscape using QGIS randomization tool. Fifteen replicas were placed within two earthen tanks while fifteen others were randomly distributed within two plots of desert scrub. To record interactions of potential predators and prey of rattlesnakes, individual replicas were tethered with fishing line approximately 2 m from a game camera (Stealth Cam, Irving, TX, USA, Stealth cam PX18FXCMO). Fishing line tethers were used to reduce any displacement by predators. Replicas were checked at least once per week to make sure that they were not displaced or broken. Cameras were set to record for 30 seconds when triggered. At each replica location, environmental data about microhabitat was collected, with microhabitat defined as a quadrat of one meter square surrounding the replica. The percentage of vegetation, exposed soil, and rock was recorded (adding up to 100%). Type of vegetation (species), vegetation height (recorded as a continuous variable), and distance to nearest vegetation (if no vegetation was within the microhabitat, continuous variable) was also recorded. Type of rock (sedimentary or conglomerate), distance to nearest rock (if no rock was within the microhabitat, continuous variable) and type of substrate (alluvial or sand) was also documented. Finally, concealment percentage, i.e. the percentage of the replica that was directly hidden from above, was recorded as a categorical variable (0%, less than 50%, or more than 50%). For each animal observed on camera, the following information was recorded: date, time, species, replica detection, and replica predation. Detection was defined as when an animal looked directly at the replica or showed any behavioral reaction to the replica (e.g. bit, pawed the replica, ran or jumped). On the other hand, predation was categorized as when a potential predator was attacking the replica with its mouth or claws. We only included records from April 1^st^ to October 31^st^ for analysis as these dates correspond to the main activity period of *Crotalus atrox* at this field site (DeSantis et al. 2019).

### Radiotracking of rattlesnakes

Twenty-five western diamond-backed rattlesnakes were caught at IMRS from 2020 to 2022 and subsequently equipped with temperature-sensitive radiotransmitters (Holohil Systems Ltd., Carp, ON, Canada, SI-2T, 9.0 g) at an on-site surgery building. Following a modified protocol derived from Hardy and Greene (2000), transmitters were implanted in the snake coelomic cavity and never exceeded 5% of the snake’s body mass. Rattlesnakes were sedated using isoflurane administered through an open-drop method within a transparent plastic tube. Benches were sterilized with isopropyl alcohol (70%), surgical instruments were sterilized in a solution of benzalkonium chloride for a minimum of 30 minutes, and procedures were conducted while wearing single-use sterile gloves. To insert the transmitter, a 1.25 cm longitudinal incision was made into the coelomic cavity, situated posteriorly at two-thirds of the snout-vent length. Subsequently, the transmitter antenna was placed subcutaneously along the body towards the head using a cannula, which was then withdrawn from the anterior, subcutaneous incision. Following the surgery, the rattlesnakes were closely observed at the on-site facility for a period of 48 hours to monitor their recovery.

Once environmental conditions were deemed favorable, rattlesnakes were released at the exact location where they had been captured. Each rattlesnake was radiotracked twice a week during one active season at IMRS (April-October 2020, 2021, 2022, and 2023). For each tracking event, the behavior of the rattlesnake was recorded as a categorical variable (tight coiled, moving, mating, non-visible). Rattlesnakes were considered active when they were visible and non-active when non-visible (usually in a burrow underground). The same microhabitat data collected for the 3D-printed replicas were collected for each tracking event (see previous description).

Animal collection was authorized by the Texas Parks and Wildlife under permit number SPR-0290-019. All animal procedures adhered to the ethical guidelines of the University of XXX and were pre-approved by the University of XXX Institutional Animal Care and Use Committee (protocol number: A-201905-2_1447328-2).

### Random forest models

Random forest models (R package *randomForest*; Liaw and Wiener 2002) were used to investigate which predictors were the most important for replica detection by predators or prey encounters, and to ultimately determine if rattlesnakes occupied space and time in a way consistent with a simple strategy of increasing prey interactions while decreasing predator encounters. Random forest models are machine learning algorithms in which classification trees are built on bootstrap samples from the data (Breiman 2001). Each tree uses a subset of randomly selected variables at each node for splitting. Out-of-bag observations, which are not included in a bootstrap sample, are used to evaluate the performance of each tree. The final prediction for each observation is determined by majority voting among the predictions of the trees, with ties being resolved randomly (Cutler et al. 2007). Variable importance can be estimated by looking at the mean decrease in accuracy. The mean decrease in accuracy for a variable is determined by comparing the classification accuracy of out-of-bag data when the variable is observed versus when the variable’s values in the out-of-bag data are randomly permuted. A higher mean decrease in accuracy signifies greater importance of the variable in classification.

To do so, game camera observations associated with their microhabitat data were used to train random forest algorithms to evaluate the importance of spatiotemporal variables for prey availability and predator detection. After training the models with this game camera dataset, the same random forest models were applied to the radiotracking dataset to estimate the probability of detection by predators or prey encounter for each tracking event. For each random forest algorithm, the number of trees was set to 500 and the best *mtry* (i.e. the number of variables randomly sampled at each split) parameter was determined based on accuracy and Kappa estimates. A summary of the methodological framework can be found in Figure 2.

**Figure 2:**
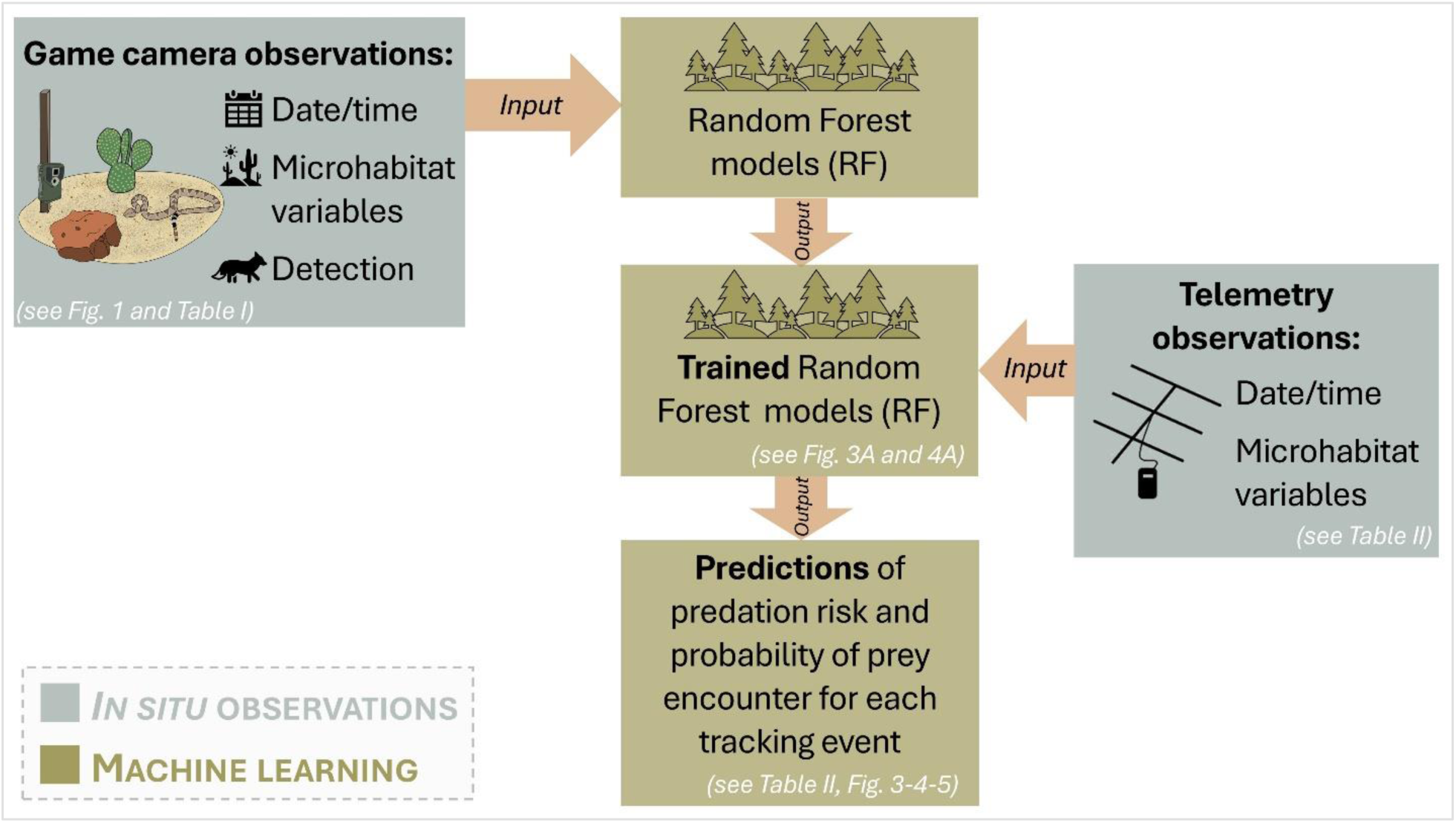
Methodological framework for the use of random forest models in this study. Results (figures and tables) are associated with each step.

### Statistical analyses

All statistical analyses were performed in R (R Core Team 2022). A *t*-test was performed on the number of recording days per camera to verify that camera trapping effort was the same between the desert scrub and earthen tank habitats. Wilcoxon signed-rank tests were used to compare the number of game camera observations of prey and predators between habitats. Wilcoxon signed-rank tests were also used to investigate the difference in detection and predation between replica sizes.

Further exploration of the predictors associated with prey encounters and detection risk was conducted using Kruskal-Wallis tests and generalized mixed models (*lme4;* Bates et al. 2015). Differences between groups were analyzed using pairwise Wilcoxon post-hoc tests. Wilcoxon signed-rank tests were used to compare the probability of detection by predators and prey encounters between tracking events where snakes were active or not (in burrows). Game camera observation data and telemetry data were used to obtain activity overlaps (Δ) between rattlesnakes and other species. Activity overlaps between organisms were calculated by measuring the similarity between the two kernel density activity curves (Ridout and Linkie 2009) using the R package *Overlap* (Meredith and Ridout 2018). Different activity overlap coefficients were calculated depending on the number of observations: Δ_1_ was used to calculate the overlap coefficient when there were less than 50 observations, while Δ_4_ was used when there were more than 50 observations (see Ridout and Linkie 2009 for formulas). To assess the significance of the activity overlap, a Mardia–Watson–Wheeler test of homogeneity was performed for each overlap coefficient (Zar 2010). Activity overlap was calculated for predators and prey for the whole active season (April-November) or divided per season: Spring (April-May), Summer (June-July), and Fall (August-October) by combining individuals from all years together (2020-2023).

## RESULTS

### Game camera observations

This study included 8,315 recording days spread across thirty game cameras. In total, the game cameras captured 5,944 observations of more than 70 species. A summary of these observations can be found in table I. Collared peccaries (*Dicotyles tajacu*), greater roadrunners (*Geococcyx californianus*) and coyotes (*Canis latrans*) represented most of the predation attempts. On the other hand, rock pocket mice (*Chaetodipus intermedius*), Merriam’s kangaroo rats (*Dipodomys merriami*), and northern mockingbird (*Mimus polyglottos*) were the most observed prey.

**Table I:** Break down of game camera observations. “Others” are species that are neither a known predator, nor prey for *C. atrox* such as mule deer (*Odocoileus hemionus*).

| Group | Number of observations | Number of species | Number of detections | Number of predation attempts |
| --- | --- | --- | --- | --- |
| Prey | 2353 | 56 | 208 | NA |
| Predator | 701 | 11 | 196 | 69 |
| Others | 2889 | 12 | 98 | NA |

### Difference between habitats and replica size

Predator observations were significantly higher in earthen tanks than in desert scrub habitats (*W* = 0; *p* < 0.001). Prey observations were also significantly higher in the earthen tanks (*W* = 21; *p* = 0.00046). Large 3D-printed replicas were significantly more detected than smaller replicas (*W* = 135; *p* = 0.03423) but the predation rate did not differ between the two (*W* = 33.5; *p* = 0.221).

### Radiotracking

In total, 936 radiotracking events from 2020 to 2022 were used for this study. Rattlesnakes were found inactive underground for 49% (n=459), tight coiled on the surface for 43% (n=405), and moving for 8% (n=72) of these observations (Table II).

**Table II:** Break down of radiotracking observations per rattlesnake behavior. The number of observations with probability of detection over 50% and with probability of prey encounter over 50% was estimated with the random forest algorithms presented in figure 2 and figure 3 respectively.

| Behavior | Number of observations | Number of observations with probability of detection over 50% | Number of observations with probability of prey encounter over 50% |
| --- | --- | --- | --- |
| Inactive | 459 | NA | NA |
| Tight coiled | 405 | 116 | 190 |
| Moving | 72 | 21 | 26 |

### Probability of detection by predators

For this random forest model, the out-of-bag accuracy was estimated at 65.15%. A breakdown of the variable importance is presented in Figure 3A. This same random forest model was then used to estimate the probability of detection for each relocation when the snake was visible (N = 490). Month had a significant effect on detection risk (*Kruskal-Wallis’ chi-squared* = 133.5, *p* < 0.001; Figure 3C) as well as time (*Kruskal-Wallis’ chi-squared* = 62.787, *p* < 0.001; Figure 3B). Concealment percentage also had a significant effect on detection risk and the probability of detection was higher when the rattlesnake was not concealed (*Kruskal-Wallis’ chi-squared* = 133.14, *p* < 0.001; Figure 3F). A summary of a generalized linear mixed model testing the effect of continuous variables on detection risk is presented in Table III.

**Figure 3:**
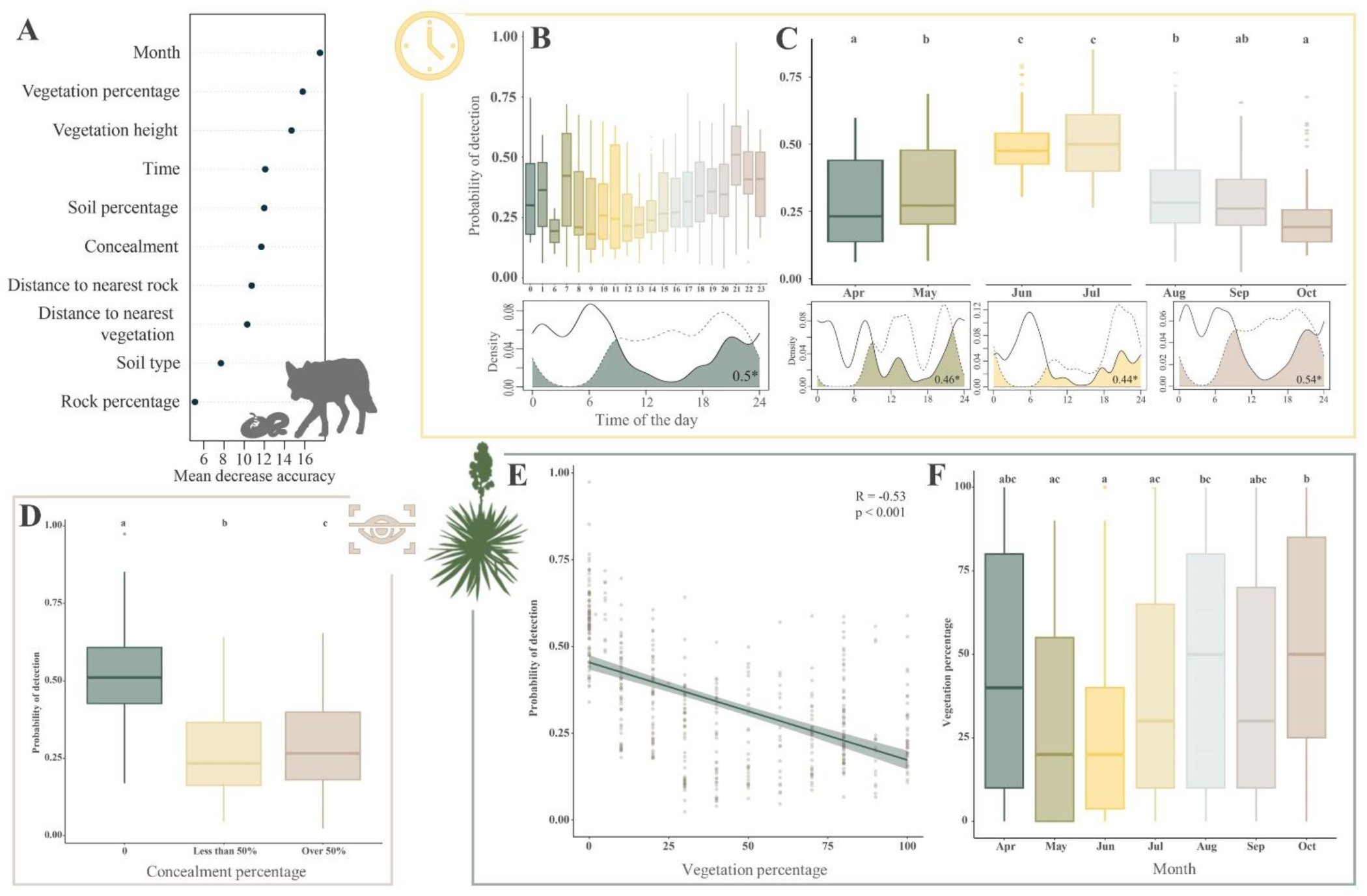
**A**: Importance of predictors associated with detection risk obtained through random forest algorithm. **B**: Box plots showing the probability of detection per hour of the day. Time of the day had a significant effect on detection risk (Kruskal-Wallis’ chi-squared, p < 0.01). Activity overlap between predators (solid line) and *Crotalus atrox* (dashed line) is represented below and the coefficient was of 0.5. **C:** Box plots showing the probability of detection per month. Month had a significant effect on detection risk (Kruskal-Wallis’ chi-squared, p < 0.01). Activity overlap between predators (solid line) and *Crotalus atrox* (dashed line) is represented below per season (April-May; June-July; Aug-Oct). Activity overlaps differed across seasons and thus, seasonal activity overlap figures have different Y-axes scales. **D:** Linear regression between the probability of detection per predator and vegetation percentage. The linear regression model (line) was significant (p < 0.001) with a coefficient of -0.53. The dots represent observations. **E:** Box plots showing vegetation percentage used per month. Month had a significant effect on detection risk (Kruskal-Wallis’ chi-squared, p < 0.01). **F:** Box plots showing the probability of detection per concealment (percentage of the snake hidden from above) category. Concealment had a significant effect on detection risk (Kruskal-Wallis’ chi-squared, p < 0.01). *Boxes indicate the inter quartile range (IQR), with the central line depicting the median and the whiskers extending to the minimal and maximal observation where outliers are represented by dots. Letters indicate the significance groups*.

**Table III:** Summary of generalized mixed model results used to investigate the effects of predictors on detection risk. SE stands for standard error. Significant p-values (<0.05) are bolded.

| Source | Estimate | SE | z | p |
| --- | --- | --- | --- | --- |
| **Probability of detection** |  |  |  |  |
| *Intercept* | 0.0308 | 0.2218 | 0.130 | 0.8896 |
| *Vegetation height* | -2.8845 | 0.5268 | -5.475 | **<0.01** |
| *Vegetation percentage* | -0.0194 | 0.0072 | -2.705 | **<0.01** |
| *Distance to nearest vegetation* | 0.6801 | 0.2039 | 3.334 | **<0.01** |

### Probability of prey encounter

The out of bag accuracy for this random forest model was estimated at 74.99%. Predictors and their respective importance are presented in Figure 3A. This model was used to obtain the probability of prey encounter for each snake relocation when the snake was visible. Time of day had a significant effect on the probability to encounter prey (*Kruskal-Wallis’ chi-squared* = 75.849, *p* < 0.001; Figure 4B). Pairwise Wilcoxon post-hoc analysis showed that month also had a significant effect on the probability of encountering prey (*Kruskal-Wallis’ chi-squared* = 146.162, *p* < 0.001; Figure 4C). Concealment had a significant effect on probability of prey encounter (*Kruskal-Wallis’ chi-squared* = 9.8948, *p* = 0.007; Figure 4E). The substrate type did not have a significant effect on the probability of prey encounter (*Kruskal-Wallis’ chi-squared* = 0.15778, *p* = 0.6912). A summary of a generalized linear mixed model testing the effect of continuous variables on probability of prey encounter is presented in Table IV.

**Figure 4:**
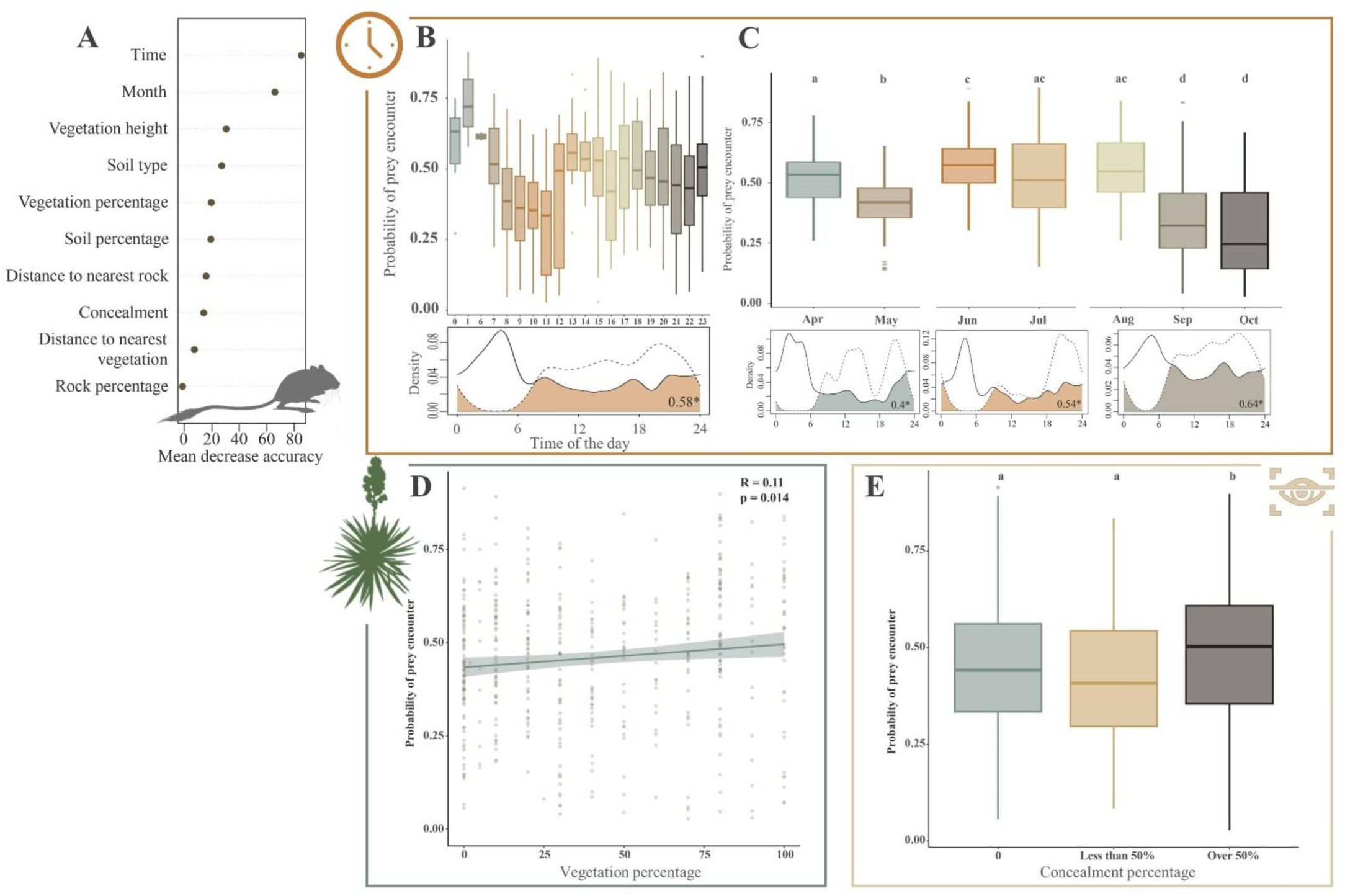
**A**: Importance of predictors associated with probability of prey encounter obtained through random forest algorithm. **B**: Box plots showing the probability of prey encounter per hour of the day. Time of the day had a significant effect on detection risk (Kruskal-Wallis’ chi-squared, p < 0.01). Activity overlap between prey (solid line) and *Crotalus atrox* (dashed line) is represented below and the coefficient was 0.58. **C:** Box plots showing the probability of prey encounter per month. Month had a significant effect on the probability of prey encounter (Kruskal-Wallis’ chi-squared, p < 0.01). Activity overlap between prey (solid line) and *Crotalus atrox* (dashed line) is represented below per season (April-May; June-July; Aug-Oct). **D:** Linear regression between the probability of prey encounter and vegetation percentage. The linear regression model (line) was significant (p = 0.014) with a coefficient of 0.11. The dots represent observations. **E:** Box plots showing the probability of detection per concealment category. Concealment had a significant effect on detection risk (Kruskal-Wallis’ chi-squared, p = 0.007). *Boxes indicate the inter quartile range (IQR), with the central line depicting the median and the whiskers extending to the minimal and maximal observation where outliers are represented by dots. Letters indicate the significance groups*.

**Table IV:** Summary of generalized mixed model results used to investigate the effects predictors on probability of prey encounter. SE stands for standard error. Significant p-values (<0.05) are bolded.

| Source | Estimate | SE | z | p |
| --- | --- | --- | --- | --- |
| **Probability of detection** |  |  |  |  |
| *Intercept* | -0.650 | 0.1742 | -3.735 | **<0.01** |
| *Vegetation height* | 0.0537 | 0.1763 | 0.305 | 0.7606 |
| *Vegetation percentage* | 0.0069 | 0.0033 | 2.113 | **0.0346** |
| *Distance to nearest rock* | 0.0017 | 0.002 | 0.847 | 0.3967 |
| *Distance to nearest vegetation* | 0.1557 | 0.1342 | 1.160 | 0.2459 |

### Activity overlap

The activity period between *Crotalus atrox* and species of interest were all significantly different from each other’s (see supplementary results for details, table S2). Overlap density plot for prey and predators are presented in figure 2 and figure 3 for the overall active seasons and per season (Spring, Summer, and Fall).

### Optimal foraging: Detection risk vs. probability of prey encounter

The probability of prey encounter (0.49) was on average significantly higher than the probability of detection by predators (0.38; *W* = 148263; *p* < 0.001). The probability of detection risk and prey encounter was compared between behavior (moving and tightly coiled) using Wilcoxon sum-rank tests (Figure 5). The probability of prey encounter was significantly higher than the probability of detection when rattlesnakes were tightly coiled on the surface (*W* = 6566, *p* <0.001). When tightly coiled, the risk of detection by predators was estimated to be over 50% for 29% of the radiotracking events while having a probability of prey encounter over 50% applied to 47% of the events (see Table II). On the other hand, no significant difference was observed when rattlesnakes were moving (*W* = 219.5, *p* = 0.1608).

**Figure 5:**
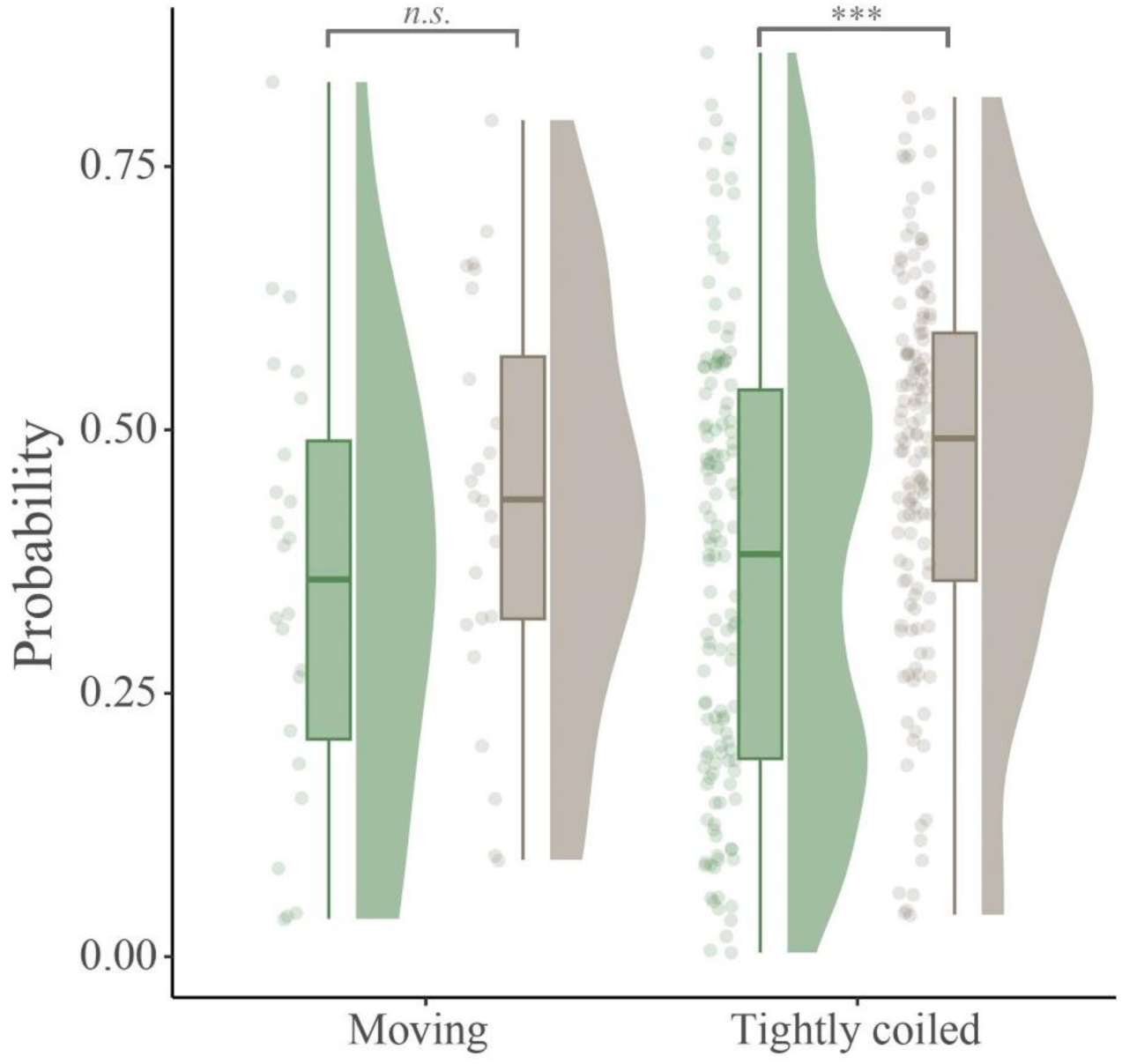
Rain cloud plots representing the probability of detection by predators (**green**) or the probability of prey encounter (**brown**) per rattlesnake behavior. Each dot represents one observation. Boxes indicate the inter quartile range with the central line depicting the median. Half violin plots show the distribution of the probability for each observation. Statistical significance between groups is denoted by ***.

## Discussion

This study aimed to determine how the landscape of fear affects the foraging decisions *of Crotalus atrox* by looking at factors influencing detection risk and prey availability using 3D-printed biologically accurate snake replicas, camera trapping, and telemetry data. Key factors with predictive capability for detection risk and prey encounter were identified using random forest models and were then applied to test hypotheses following optimal foraging theory. Rattlesnakes overlapped both spatially and temporally with prey and predators, but on average, rattlesnakes had significantly higher probability of prey encounter than predator detection, supporting optimal foraging theory. This study also highlighted the effectiveness of using 3D-printed snake replicas paired with game cameras and associated with telemetry data and machine learning to test hypotheses of predator-prey interactions. This valuable new combination of tools helped further our understanding of cryptic mesopredators, especially in a context of predator-prey interactions.

The probability of detection by predators varied temporally throughout the active season (April-October). Detection probability fluctuated daily with higher probability of detection by predators at night from 21:00 to 23:00 hours. This timeframe corresponds partially to the time interval in which rattlesnakes and their predators were the most active. Predators were mostly active from 20:00 to 06:00 hours which was also observed in similar studies conducted in the Chihuahuan Desert (Bissonette 1978; Durán-Antonio et al. 2020). Rattlesnakes were mostly active from 18:00 to 23:00. The activity period of rattlesnakes and their predators were significantly different suggesting that rattlesnakes temporally avoid predators as a strategy to decrease their risk of predation (Creel 2018). However, the diel activity pattern of rattlesnakes might be skewed in this study due to the lack of sampling effort between 01:00 and 06:00 hours. Indeed, *C. atrox* has been reported to mostly move between 18:00 and 06:00 hours at the same study site (DeSantis et al. 2020). Rattlesnakes might also be more active during the day when their predators are not due to their thermal requirements. Collared peccaries and coyotes tend to avoid being active during the day and switch to a more nocturnal lifestyle when the temperatures rise above a certain threshold (Bissonette 1978; Melville et al. 2020). Rattlesnake activity is directly correlated to ambient temperature with the proportion of time hunting on the surface decreasing when average daily air temperatures are increasing (Putman and Clark 2015). Nevertheless, rattlesnakes also need to engage in basking at certain times of the year during the day when their predators might not be active, to attain a specific temperature to facilitate essential physiological processes (Huey 1982). Even if a significant difference in activity periods between rattlesnakes and their predators was maintained throughout the year, predator detection varied significantly per month with higher probability to be detected in June and July. These results were not explained by the number of predator detections observed or the number of rattlesnakes observations per month. However, the vegetative structure used by rattlesnakes, including both microhabitat percentage and vegetation height, exhibited significant monthly variations with June recording the lowest percentage and height, and April and May the highest. Conversely, the probability of detection significantly increased when percentage of vegetation decreased, meaning that variations in vegetative cover could explain the monthly difference in detection probability. Vegetation provides concealment to rattlesnakes, which was also linked to a decrease in probability of being detected in this study. Habitat structure and complexity have been linked to predation risk in other studies (Denno et al. 2005; Gigliotti et al. 2020; Duchesne et al. 2022). Complex habitat structure provides shelter for organisms (Stellatelli et al. 2015; Worthington-Hill and Gill 2019), hinders the movements of predators (Ferreira et al. 2018), and decreases the visual detection of prey (Allen et al. 2013; Law et al. 2020), resulting in an overall decrease in predation success.

The probability to encounter prey also varied temporarily with a significant effect of time of the day and month. *Crotalus atrox* are known to prey upon rodents, birds, and lizards (Beavers 1976). In this study, rodents, mostly represented by rock pocket mice (*Chaetodipus intermedius*) and Merriam’s Kangaroo rats (*Dipodomys merriami*), exhibited a nocturnal activity period. The activity period of passerine birds followed a bimodal distribution with peaks around sunrise and sunset. *Crotalus atrox* activity followed the same bimodal distribution as birds during the spring and the summer. It is unlikely that *Crotalus atrox* matches its activity pattern to those of birds as birds only constitute only a small portion of their diet (Beavers 1976). Many species of birds are active around sunrise and sunset to reduce predation risk (Bednekoff and Houston 1994; McNamara et al. 1994; Reyes-Arriagada et al. 2015) but also to avoid high temperatures (Silva et al. 2015). *Crotalus atrox* likely shows the same bimodal distribution to also avoid high temperature in the middle of the day. The activity overlap between rattlesnakes and rodents is expected to be much higher during the night as rattlesnakes are known to be active throughout the night (DeSantis et al. 2020), but sampling effort was limited between 01:00 and 06:00 hours. The month of the year also had a significant effect on the probability of encountering prey, which did not appear to be caused by seasonal changes in activity overlap. Probability of prey encounter was smaller in September and October, which corresponds mostly to the mating season of *Crotalus atrox* at this field site (DeSantis et al. 2019). Because of their ambush foraging strategy, male rattlesnakes face a trade-off between hunting and searching for mates (Tetzlaff et al. 2017). Consequently, male snakes often decrease their foraging behavior (Slip and Shine 1988; Daltry et al. 1998; Glaudas and Alexander 2016; Tetzlaff et al. 2017; Cochran et al. 2021) to increase their reproductive output (Tetzlaff et al. 2017), as observed in this study. Vegetation percentage also had a positive significant effect on probability of prey encounter. Vegetation structure and community are predictors of both rodent (Kluever et al. 2016) and bird abundance (Macías-Duarte et al. 2018). Vegetation also provides concealment from predators (as shown in this study) and thus, might offer shelter and foraging opportunities for rodents and birds. In fact, concealment had a significant effect on the probability of encountering prey, with a higher chance when rattlesnakes are 50% or more concealed. Rattlesnakes may be taking advantage of concealment to mediate predation risk and concurrently increase their foraging opportunities when active on the surface.

Rattlesnakes appeared to be active where and when both predators and prey were also active. However, the probability of encountering prey for rattlesnakes was significantly higher than being detected by a predator. For the 405 relocations where rattlesnakes were tightly coiled on the surface, rattlesnakes had a 50% chance or more to encounter prey for 190 of these events (47%), whereas they had a 50% chance or more to be detected by a predator for only 116 of these events (29%). This result could be explained by the higher density of prey compared to predators at our field site, or because rattlesnakes select less risky microhabitat with higher chance to encounter prey, thus following optimal foraging theory. When comparing the probability of detection by predators and the probability of prey encounter, rattlesnake behavior was important. The probability of prey encounter was significantly higher than detection risk when rattlesnakes were tightly coiled on the surface, but not when they were moving. When rattlesnakes are tightly coiled on the surface, they are generally in ambush waiting for prey or thermoregulating. In this study, rattlesnakes chose to be tightly coiled when and where the probability of encountering prey was significantly higher than the probability of detection by predators. Because of the cryptic nature of rattlesnakes (Da Cunha et al. 2024), the predation pressure exerted on them appears to be relatively low when they are tightly coiled on the surface (Maag and Clark 2022). Instead, predation likely happens when rattlesnakes are performing risky behavior such as moving (Maag and Clark 2022). However, the effect of movement could not be considered in this study as the 3D-printed replicas, despite mimicking a moving rattlesnake, were static.

3D-printed snake replicas were suitable to study predation in situ. These replicas resisted the environmental conditions of the Chihuahuan Desert for two years as well as the mechanical pressures exerted by the jaws of collared peccaries, horses, and coyotes. 3D-printed replicas offered an efficient alternative to the traditional plasticine replicas (Behm et al. 2018; Bulté et al. 2018) as they did not require frequent checks for predation marks while enabling the identification of predators down to the species level when paired with a game-camera. These 3D-printed replicas were successfully used to determine the main predators of *Crotalus atrox* at this field site: collared peccaries, greater roadrunners, and coyotes. Greater roadrunners and coyotes have been previously reported as predators of rattlesnakes (Klauber 1956*b*; Maag and Clark 2022). Surprisingly, collared peccaries accounted for most of the predation attempts. When encountering a rattling rattlesnake, collared peccaries tend to withdraw quickly, but are known to kill snakes (Neal 1959). While collared peccaries may not primarily eat rattlesnakes, they may occasionally consume them, particularly if the snakes are found dead. Thus, the high number of recorded instances of predation by collared peccaries on the 3D replicas could be attributed to the possibility that the peccaries perceived the rattlesnake replicas as lifeless, given their static nature. The combination of 3D-printed replicas and game cameras proved to be effective not only in identifying species that predate *Crotalus atrox* but also in determining the primary factors influencing predation and predator detection rates. Overall, the detection rates for all animals were affected by the size of the 3D-printed replicas, with larger replicas being detected more frequently than smaller ones. However, the predation rate between small and large replicas did not significantly differ. These finding could be attributed to the fact that the primary predators identified (collared peccaries, coyotes) were sufficiently large to kill and prey upon adult rattlesnakes found in West Texas (Klauber 1956*a*; Neal 1959).

In conclusion, this study provided new insights on how mesopredators navigate the landscape of fear to optimize foraging by demonstrating the interplay between predator detection risk and probability of prey encounter. Indeed, rattlesnakes appeared to choose to be active when and where the probability of prey encounter was significantly higher than the risk of predation. Future research should explore additional factors, including meteorological data and specific behaviors such as movement, to further refine our comprehension of rattlesnake foraging strategy. By combining tools such as 3D-printed biologically accurate replicas, camera trapping, telemetry, and machine learning models, this study contributed to the understanding of optimal foraging theory while validating a novel approach to study cryptic mesopredator foraging decisions. By identifying the key factors influencing predation risk and prey encounter, this study not only furthers our understanding of predator-prey interactions but also provides a framework for future studies investigating optimal foraging theory in natural settings.

## Supporting information

Supplementary Materials and Results

## Literature cited

1. Allen, W. L., R. Baddeley, N. E. Scott-Samuel, and I. C. Cuthill. 2013. The evolution and function of pattern diversity in snakes. Behavioral Ecology 24:1237–1250.

2. Bates, D., M. Mächler, B. Bolker, and S. Walker. 2015. Fitting Linear Mixed-Effects Models Using lme4 67:1–48.

3. Beavers, R. A. 1976. Food Habits of the Western Diamondback Rattlesnake, Crotalus atrox, in Texas (Viperidae). The Southwestern Naturalist 20:503–515.

4. Bednekoff, P. A., and A. I. Houston. 1994. Avian daily foraging patterns: Effects of digestive constraints and variability. Evolutionary Ecology 8:36–52.

5. Behm, J. E., B. R. Waite, S. T. Hsieh, and M. R. Helmus. 2018. Benefits and limitations of three-dimensional printing technology for ecological research. BMC Ecology 18:32.

6. Belgrad, B. A., and B. D. Griffen. 2016. Predator–prey interactions mediated by prey personality and predator hunting mode. Proceedings of the Royal Society B: Biological Sciences 283:20160408.

7. Berger, J., P. B. Stacey, L. Bellis, and M. P. Johnson. 2001. A Mammalian Predator–Prey Imbalance: Grizzly Bear and Wolf Extinction Affect Avian Neotropical Migrants. Ecological Applications 11:947– 960.

8. Bissonette, J. A. 1978. The Influence of Extremes of Temperature on Activity Patterns of Peccaries. The Southwestern Naturalist 23:339–346.

9. Bleicher, S. S. 2017. The landscape of fear conceptual framework: definition and review of current applications and misuses. PeerJ 5:e3772.

10. Breiman, L. 2001. Random Forests. Machine Learning 45:5–32.

11. Brown, J. S., and B. P. Kotler. 2004. Hazardous duty pay and the foraging cost of predation. Ecology Letters 7:999–1014.

12. Bulté, G., R. J. Chlebak, J. W. Dawson, and G. Blouin-Demers. 2018. Studying mate choice in the wild using 3D printed decoys and action cameras: a case of study of male choice in the northern map turtle. Animal Behaviour 138:141–143.

13. Cochran, C., K. L. Edwards, Z. D. Travis, L. R. Pompe, and W. K. Hayes. 2021. Diet and Feeding Frequency in the Southwestern Speckled Rattlesnake (Crotalus pyrrhus): Ontogenetic, Sexual, Geographic, and Seasonal Variation. Journal of Herpetology 55:77–87.

14. Creel, S. 2018. The control of risk hypothesis: reactive vs. proactive antipredator responses and stress-mediated vs. food-mediated costs of response. Ecology Letters 21:947–956.

15. Creel, S., J. Winnie Jr., B. Maxwell, K. Hamlin, and M. Creel. 2005. Elk Alter Habitat Selection as an Antipredator Response to Wolves. Ecology 86:3387–3397.

16. Cresswell, W. 2011. Predation in bird populations. Journal of Ornithology 152:251–263.

17. Cutler, D. R., T. C. Edwards Jr., K. H. Beard, A. Cutler, K. T. Hess, J. Gibson, and J. J. Lawler. 2007. Random Forests for Classification in Ecology. Ecology 88:2783–2792.

18. Da Cunha, O., C. Fournier, L. M. Horne, B. M. Seymoure, and J. D. Johnson. 2024. You can’t see me: Background matching in the Western diamond-backed rattlesnake (Crotalus atrox) J. Zool.

19. Daltry, J. C., W. Wüster, and R. S. Thorpe. 1998. Intraspecific Variation in the Feeding Ecology of the Crotaline Snake Calloselasma rhodostoma in Southeast Asia. Journal of Herpetology 32:198–205.

20. Denno, R. F., D. L. Finke, and G. A. Langelloto. 2005. Direct and indirect effects of vegetation structure and habitat complexity on predator-prey and predator-predator interactions. Pages 211–239 in Ecology of predator-prey interactions. Oxford University Press, Oxford, UK.

21. DeSantis, D. L., V. Mata-Silva, J. D. Johnson, and A. E. Wagler. 2020. Integrative Framework for Long-Term Activity Monitoring of Small and Secretive Animals: Validation With a Cryptic Pitviper. Frontiers in Ecology and Evolution 8.

22. DeSantis, D. L., A. E. Wagler, V. Mata-Silva, and J. D. Johnson. 2019. Effects of human-made resource hotspots on seasonal spatial strategies by a desert pitviper. Scientific Reports 9:16690.

23. Duchesne, T., E. Graitson, O. Lourdais, S. Ursenbacher, and M. Dufrêne. 2022. Fine-scale vegetation complexity and habitat structure influence predation pressure on a declining snake. Journal of Zoology 318:205–217.

24. Durán-Antonio, J., A. González-Romero, and V. J. Sosa. 2020. Activity overlap of carnivores, their potential wild prey, and temporal segregation, with livestock in a Biosphere Reserve in the Chihuahuan Desert. Journal of Mammalogy 101:1609–1621.

25. Emlen, J. M. 1966. The Role of Time and Energy in Food Preference. The American Naturalist 100:611– 617.

26. Farallo, V. R., and M. R. J. Forstner. 2012. Predation and the Maintenance of Color Polymorphism in a Habitat Specialist Squamate. PLOS ONE 7:e30316.

27. Ferreira, A. S., C. A. Peres, J. A. Bogoni, and C. R. Cassano. 2018. Use of agroecosystem matrix habitats by mammalian carnivores (Carnivora): a global-scale analysis. Mammal Review 48:312–327.

28. Fortin, D., H. L. Beyer, M. S. Boyce, D. W. Smith, T. Duchesne, and J. S. Mao. 2005. Wolves Influence Elk Movements: Behavior Shapes a Trophic Cascade in Yellowstone National Park. Ecology 86:1320– 1330.

29. Gaynor, K. M., J. S. Brown, A. D. Middleton, M. E. Power, and J. S. Brashares. 2019. Landscapes of Fear: Spatial Patterns of Risk Perception and Response. Trends in Ecology & Evolution 34:355–368.

30. Gigliotti, L. C., R. Slotow, L. T. B. Hunter, J. Fattebert, C. Sholto-Douglas, and D. S. Jachowski. 2020. Habitat complexity and lifetime predation risk influence mesopredator survival in a multi-predator system. Scientific Reports 10:17841.

31. Glaudas, X., and G. J. Alexander. 2016. Food supplementation affects the foraging ecology of a low-energy, ambush-foraging snake. Behavioral Ecology and Sociobiology 71:5.

32. Grabowski, J. H., and D. L. Kimbro. 2005. Predator-Avoidance Behavior Extends Trophic Cascades to Refuge Habitats. Ecology 86:1312–1319.

33. Hardy, D. L., and H. W. Greene. 2000. Inhalation anesthesia of rattlesnakes in the field for processing and transmitter implantation 13:109–113.

34. Harmel, M. V., H. L. Crowell, J. M. Whelan, and E. N. Taylor. 2020. Rattlesnake colouration affects detection by predators. Journal of Zoology 311:260–268.

35. Haswell, P. M., K. A. Jones, J. Kusak, and M. W. Hayward. 2018. Fear, foraging and olfaction: how mesopredators avoid costly interactions with apex predators. Oecologia 187:573–583.

36. Hawlena, D., and O. J. Schmitz. 2010. Physiological Stress as a Fundamental Mechanism Linking Predation to Ecosystem Functioning. The American Naturalist 176:537–556.

37. Huey, R. B. 1982. Temperature, physiology, and the ecology of reptiles. Physiological ecology 25–95.

38. Igic, B., V. Nunez, H. U. Voss, R. Croston, Z. Aidala, A. V. López, A. V. Tatenhove, et al. 2015. Using 3D printed eggs to examine the egg-rejection behaviour of wild birds. PeerJ 3:e965.

39. Kishida, O., Z. Costa, A. Tezuka, and H. Michimae. 2014. Inducible offences affect predator–prey interactions and life-history plasticity in both predators and prey. Journal of Animal Ecology 83:899–906.

40. Klauber, L. M. 1956a. Rattlesnakes: their habits, life histories, and influence on mankind. University of California Press, Berkeley.

41. Kishida, O., Z. Costa, A. Tezuka, and H. Michimae. 1956b. Rattlesnakes (Vol. 1). University of California Press, Berkeley.

42. Kluever, B. M., E. M. Gese, and S. J. Dempsey. 2016. The influence of wildlife water developments and vegetation on rodent abundance in the Great Basin Desert. Journal of Mammalogy 97:1209–1218.

43. Kneitel, J. M., and J. M. Chase. 2004. Disturbance, Predator, and Resource Interactions Alter Container Community Composition. Ecology 85:2088–2093.

44. Law, C., L. Lancaster, J. Hall, S. Handy, M. Hinchliffe, C. O’Brien, K. O’Brien, et al. 2020. Quantifying the differences in avian attack rates on reptilesbetween an infrastructure and a control site. European Journal of Wildlife Research 66:54.

45. Letnic, M., E. G. Ritchie, and C. R. Dickman. 2012. Top predators as biodiversity regulators: the dingo Canis lupus dingo as a case study. Biological Reviews 87:390–413.

46. Liaw, A., and M. Wiener. 2002. Classification and Regression by randomForest 2.

47. Lima, S. L. 1998. Nonlethal Effects in the Ecology of Predator-Prey Interactions. BioScience 48:25–34.

48. Lima, S. L. 2009. Predators and the breeding bird: behavioral and reproductive flexibility under the risk of predation. Biological Reviews 84:485–513.

49. Maag, D., and R. Clark. 2022. Safety in coils: predation rates of ambush hunting rattlesnakes are extremely low. Amphibia-Reptilia 43:425–430.

50. MacArthur, R. H., and E. R. Pianka. 1966. On Optimal Use of a Patchy Environment. The American Naturalist 100:603–609.

51. Macías-Duarte, A., A. O. Panjabi, D. B. Pool, I. Ruvalcaba-Ortega, and G. J. Levandoski. 2018. Fall vegetative cover and summer precipitation predict abundance of wintering grassland birds across the Chihuahuan desert. Journal of Arid Environments 156:41–49.

52. McGhee, K. E., L. M. Pintor, and A. M. Bell. 2013. Reciprocal Behavioral Plasticity and Behavioral Types during Predator-Prey Interactions. The American Naturalist 182:704–717.

53. McNamara, J. M., A. I. Houston, and S. L. Lima. 1994. Foraging Routines of Small Birds in Winter: A Theoretical Investigation. Journal of Avian Biology 25:287–302.

54. Melville, H. I. A. S., W. C. Conway, J. B. Hardin, C. E. Comer, and M. L. Morrison. 2020. Abiotic variables influencing the nocturnal movements of bobcats and coyotes. Wildlife Biology 2020:wlb.00601.

55. Meredith, M., and M. Ridout. 2018. Overview of the overlap package. R Project.

56. Mooney, K. A., D. S. Gruner, N. A. Barber, S. A. Van Bael, S. M. Philpott, and R. Greenberg. 2010. Interactions among predators and the cascading effects of vertebrate insectivores on arthropod communities and plants. Proceedings of the National Academy of Sciences 107:7335–7340.

57. Mukherjee, S., M. Zelcer, and B. P. Kotler. 2009. Patch use in time and space for a meso-predator in a risky world. Oecologia 159:661–668.

58. Neal, B. J. 1959. A Contribution on the Life History of the Collared Peccary in Arizona. The American Midland Naturalist 61:177–190.

59. Nelson, E. H., C. E. Matthews, and J. A. Rosenheim. 2004. Predators Reduce Prey Population Growth by Inducing Changes in Prey Behavior. Ecology 85:1853–1858.

60. Pangle, K. L., S. D. Peacor, and O. E. Johannsson. 2007. Large Nonlethal Effects of an Invasive Invertebrate Predator on Zooplankton Population Growth Rate. Ecology 88:402–412.

61. Peckarsky, B. L., C. A. Cowan, M. A. Penton, and C. Anderson. 1993. Sublethal Consequences of Stream-Dwelling Predatory Stoneflies on Mayfly Growth and Fecundity. Ecology 74:1836–1846.

62. Pettorelli, N., A. Hilborn, C. Duncan, and S. M. Durant. 2015. Chapter Two - Individual Variability: The Missing Component to Our Understanding of Predator–Prey Interactions. Pages 19–44 in S. Pawar, G. Woodward, and A. I. Dell, eds. Advances in Ecological Research, Trait-Based Ecology - From Structure to Function (Vol. 52). Academic Press.

63. Putman, B. J., and R. W. Clark. 2015. Habitat Manipulation in Hunting Rattlesnakes (Crotalus Species). The Southwestern Naturalist 60:374–377.

64. Pyke, G. H., H. R. Pulliam, and E. L. Charnov. 1977. Optimal Foraging: A Selective Review of Theory and Tests. The Quarterly Review of Biology 52:137–154.

65. R Core Team. 2022. R: A Language and Environment for Statistical Computing. R Foundation for Statistical Computing, Vienna, Austria.

66. Reyes-Arriagada, R., J. E. Jiménez, and R. Rozzi. 2015. Daily patterns of activity of passerine birds in a Magellanic sub-Antarctic forest at Omora Park (55°S), Cape Horn Biosphere Reserve, Chile. Polar Biology 38:401–411.

67. Ridout, M. S., and M. Linkie. 2009. Estimating overlap of daily activity patterns from camera trap data. Journal of Agricultural, Biological, and Environmental Statistics 14:322–337.

68. Schneider, C. A., W. S. Rasband, and K. W. Eliceiri. 2012. NIH Image to ImageJ: 25 years of image analysis. Nature Methods 9:671–675.

69. Silva, J. P., I. Catry, J. M. Palmeirim, and F. Moreira. 2015. Freezing heat: thermally imposed constraints on the daily activity patterns of a free-ranging grassland bird. Ecosphere 6:art119.

70. Skelly, D. K., and E. E. Werner. 1990. Behavioral and Life-Historical Responses of Larval American Toads to an Odonate Predator. Ecology 71:2313–2322.

71. Slip, D. J., and R. Shine. 1988. Feeding Habits of the Diamond Python, Morelia s. spilota: Ambush Predation by a Boid Snake. Journal of Herpetology 22:323–330.

72. Stellatelli, O. A., C. Block, L. E. Vega, and F. B. Cruz. 2015. Nonnative Vegetation Induces Changes in Predation Pressure and Escape Behavior of Two Sand Lizards (Liolaemidae: Liolaemus). Herpetologica 71:136–142.

73. Tetzlaff, S. J., E. T. Carter, B. A. DeGregorio, M. J. Ravesi, and B. A. Kingsbury. 2017. To forage, mate, or thermoregulate: Influence of resource manipulation on male rattlesnake behavior. Ecology and Evolution 7:6606–6613.

74. Troscianko, J., and M. Stevens. 2015. Image calibration and analysis toolbox – a free software suite for objectively measuring reflectance, colour and pattern. Methods in Ecology and Evolution 6:1320–1331.

75. Wasserman, R. J., M. E. Alexander, O. L. F. Weyl, D. Barrios-O’Neill, P. W. Froneman, and T. Dalu. 2016. Emergent effects of structural complexity and temperature on predator–prey interactions. Ecosphere 7:e01239.

76. Winnie, J., and S. Creel. 2007. Sex-specific behavioural responses of elk to spatial and temporal variation in the threat of wolf predation. Animal Behaviour 73:215–225.

77. Worthington, R. D., J. D. Johnson, C. S. Lieb, and W. Anderson. 2022. Biotic Resources of Indio Mountains Research Station (IMRS) Southeastern Hudspeth County, Texas: A Handbook for Students and Researchers. University of Texas at El Paso, USA.

78. Worthington-Hill, J. O., and J. A. Gill. 2019. Effects of large-scale heathland management on thermal regimes and predation on adders Vipera berus. Animal Conservation 22:481–492.

79. Zar, J. H. 2010. Biostatistical analysis (5th edition.). Pearson Education, Inc., Hoboken, New Jersey.

