## Supplementary Materials and Results for "From Fear to Feast: Rattlesnakes Navigate the Landscape of Fear to Optimize Foraging"

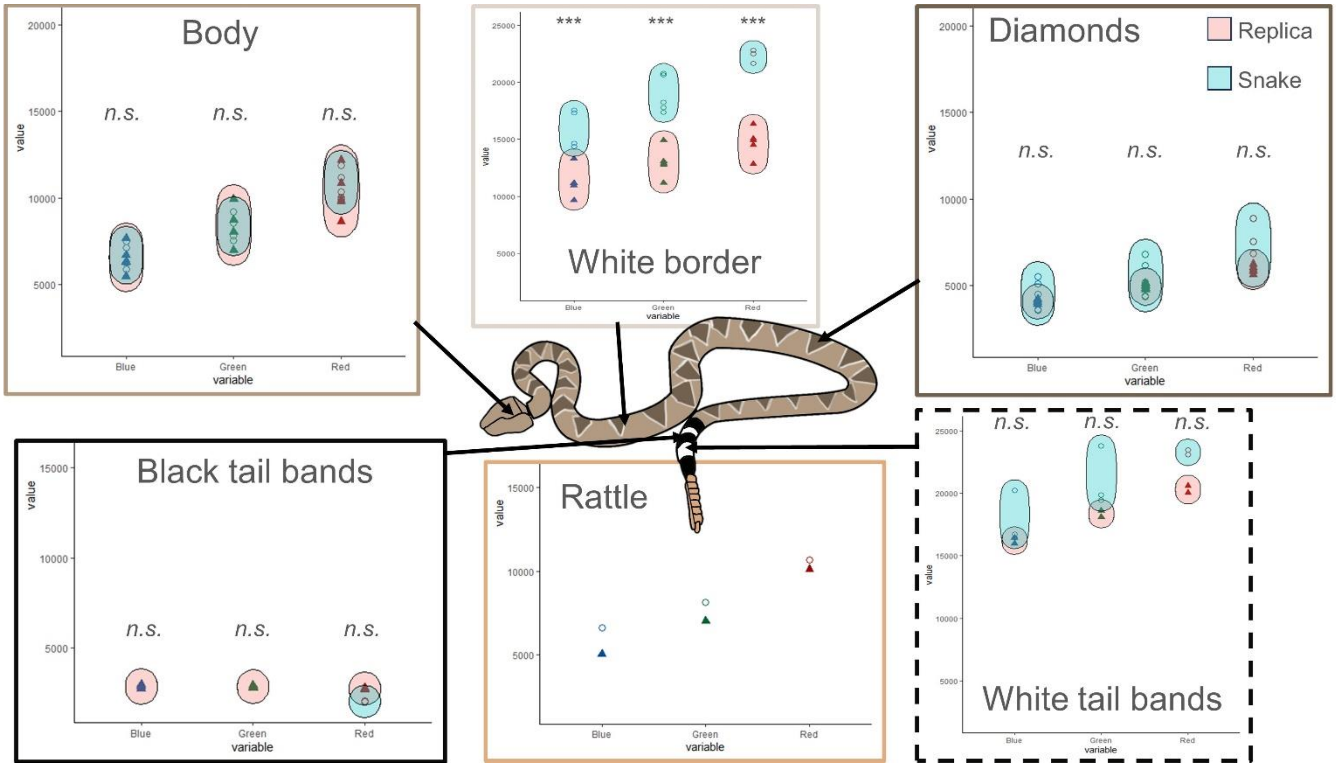


**Figure S1:** Color measurement of different body regions from a live specimen of *Crotalus atrox* (blue) and a 3D-printed replica (red). The value on the y-axis corresponds to the mean pixel value of the area measured in the blue, green, and red channel. Paint coloration and body coloration were significantly matching for all body parts, beside the white border found around the diamonds.

**Table S1:** Specific paints and mixes used to paint the replicas. Paints were from the brand Acrylicos Vallejo. Units are in cubic centimeters (cc).

| Body part | Paint mix |
|---|---|
| Body primer | 1.5cc *White* + 0.5cc *Light Gray* + 2cc distilled water |
| Body | 3cc *Iraqi Sand* + 1cc *Light Gray* + 1cc *Leather Brown* + 5cc distilled water |
| Diamonds | 1cc *Iraqi Sand* + 1cc *Flat Earth* + 3cc *Leather Brown* + 5cc distilled water |
| Rattle | 3cc *Iraqi Sand* + 1cc *Flat Earth* + 4cc distilled water |
| White border | 1.5cc *White* + 0.5cc *Iraqi Sand* + 2cc distilled water |
| Black tail bands | 3cc *Iraqi Sand* + 1cc *Black* + 4cc distilled water |
| White tail bands | 3cc *White* + 3cc distilled water |

### SUPPLEMENTARY RESULTS

**Table S2:** Summary of overlap coefficient and their significance (according to Mardia-Watson-Wheeler test of homogeneity) for predators and prey with *Crotalus atrox*. Overlap could not be calculated between *G. califonianus* and *C. atrox* for the spring as only one individual *G. califonianus* was observed.

| Species | Season | Coefficient | Significance |
|---|---|---|---|
| All predators | Year | 0.5 | 2.20E-16 |
| *C. latrans* | Year | 0.55 | 2.20E-16 |
| *D. tajacu* | Year | 0.44 | 2.20E-16 |
| *G. californianus* | Year | 0.52 | 1.29E-06 |
| All predators | Spring | 0.46 | 3.61E-07 |
| *C. latrans* | Spring | 0.49 | 0.0007822 |
| *D. tajacu* | Spring | 0.49 | 0.01228 |
| *G. californianus* | Spring | *N.A.* | *N.A.* |
| All predators | Summer | 0.44 | 1.43E-14 |
| *C. latrans* | Summer | 0.41 | 1.09E-06 |
| *D. tajacu* | Summer | 0.39 | 1.86E-13 |
| *G. californianus* | Summer | 0.32 | 0.0001939 |
| All predators | Fall | 0.54 | 2.20E-16 |
| *C. latrans* | Fall | 0.57 | 2.20E-16 |
| *D. tajacu* | Fall | 0.45 | 2.20E-16 |
| *G. californianus* | Fall | 0.57 | 0.0006186 |
| All prey | Year | 0.58 | 2.20E-16 |
| Birds | Year | 0.712 | 0.0002 |
| Rodents | Year | 0.36 | 2.20E-16 |
| All prey | Spring | 0.4 | 1.31E-10 |
| Birds | Spring | 0.77 | 4.08E-02 |
| Rodents | Spring | 0.19 | 2.20E-16 |
| All prey | Summer | 0.54 | 2.20E-12 |
| Birds | Summer | 0.55 | 5.44E-12 |
| Rodents | Summer | 0.34 | 2.20E-16 |
| All prey | Fall | 0.64 | 2.20E-16 |
| Birds | Fall | 0.72 | 1.53E-02 |
| Rodents | Fall | 0.33 | 2.20E-16 |
